# Tandem mass spectrum similarity-based network analysis using ^13^C-labeled and non-labeled metabolome data to identify the biosynthesis pathway of the blood pressure-lowering asparagus metabolite asparaptine A

**DOI:** 10.1101/2021.02.09.430543

**Authors:** Ryo Nakabayashi, Yutaka Yamada, Tomoko Nishizawa, Tetsuya Mori, Takashi Asano, Masanari Kuwabara, Kazuki Saito

## Abstract

Asparaptine, a conjugate of arginine and asparagusic acid, was found in asparagus (*Asparagus officinalis*) as a naturally occurring inhibitor of angiotensin-converting enzyme (ACE) *in vitro.* The biosynthetic pathway to asparaptine is largely unknown; however, it is suggested that asparagusic acid may be biosynthesized from valine. To determine which metabolites are involved in the asparaptine biosynthetic pathway, we performed tandem mass spectrometry similarity-based metabolome network analysis using ^13^C labeled and non-labeled valine-fed asparagus calluses. We determined that valine is used as a starting material, *S*(2-carboxy-*n*-propyl)-cysteine as an intermediate, and two new metabolites as asparaptine analogs, lysine- and histidine-type conjugates, are involved in the pathway. Asparaptine was therefore renamed asparaptine A (arginine type), and the two analogs were named asparaptines B (lysine type) and C (histidine type). Oral feeding of asparaptine A to a hypertensive mouse species showed that this metabolite lowers both blood pressure and heart rate within two hours and both of which were back to normal two days later. These results suggest that asparaptine A may not only have effects as an ACE inhibitor, but also has β-antagonistic effects, which are well-known to be preventive for cardiovascular diseases.

## Introduction

Stable isotope labeling is a powerful approach in metabolomics for determining the number of elements in detected metabolites using liquid chromatography-tandem mass spectrometry (LC-MS/MS)^1^. LC-MS/MS analysis is useful for narrowing down metabolite identities using peak resolution and mass accuracy^2^, but it is difficult to determine unambiguous molecular formula. A metabolite labeled with stable isotope (^13^C, ^15^N, ^18^O, or ^34^S) and its non-labeled counterpart are typically detected at almost the same retention time and can be paired. The shifted mass to charge (*m/z*) value between metabolite pairs allows the numbers of C, N, O, or S to be determined with an established procedure in plants^3–9^. Recently, principal component analysis (PCA) was used for pairing fully ^15^N-labeled and non-labeled metabolites to characterize the missing monoterpene indole alkaloids in *Catharanthus roseus*^10^. The PCA clearly separated types of metabolites that differed significantly between samples. In that study, pairing was manually performed using coordinates of loading factors in the PCA as well as retention times and *m/z* values of the precursor ions. Manual pairing approaches are suitable for chemical assignments of dozens of metabolites with the caveat of requiring long data analysis time; however, are unfeasible for assignments of metabolome that can include hundreds of metabolites.

To streamline the processes involved, it is essential to apply methods of computational automation. For instance, molecular networking by Global Natural Product Society^11,12^ or MS-DIAL/FINDER^3,13^ programs enable the creation of MS/MS similarity-based networks that can assist structural analog characterization. Theoretically, the fragmentation “pattern” of isotope-labeled and non-labeled metabolites should be almost identical since the chemical properties of the metabolites, apart from their *m/z* value, are otherwise the same. It is possible to pair stable isotope-labeled and non-labeled MS/MS spectra by using MS/MS similarity scores in addition to retention times and *m/z* values.

Asparaptine, which consists of arginine and asparagusic acid, is a naturally occurring inhibitor of angiotensin-converting enzyme (ACE) in asparagus (*Asparagus officinalis*)^14^. It has been suggested that the asparagusyl moiety of asparaptine is biosynthesized from valine^15^; however, the biosynthetic pathway to asparagusic acid and asparaptine remains largely unknown. In this study, we performed MS/MS spectrum similarity-based network analysis using ^13^C-labeled and non-labeled metabolome data to determine metabolites involved in the biosynthetic pathway of asparaptine. We showed that valine is a starting material for the biosynthesis of asparaptine and that *S*-(2-carboxy-*n*-propyl)-L-cysteine is an intermediate of the pathway. Moreover, we found two new structural analogs of asparaptine: conjugates of lysine/histidine and asparagusic acid. Thus, asparaptine was renamed as asparaptine A, and the lysine- and histidine-type conjugates were named as asparaptine B and C, respectively. Finally, we characterized that asparaptine A lowers blood pressure and heart rate of a hypertensive mouse species.

## Experimental Section

### Plant materials

Calluses derived from green asparagus (*Asparagus officinalis*)^16^ were used in this study.

### Chemicals

The following ^13^C amino acids and non-labeled amino acids were used in the study: L-valine (^13^C5, 99%), L-lysine (^13^C6, 99%), L-histidine (^13^C6, 97%-99%), and L-glutamine (^13^C5, 99%), (Cambridge Isotope Laboratories, US); and L-valine, L-lysine, and glutamine (Sigma Aldrich Japan, Tokyo), and L-arginine and L-histidine (FUJIFILM Wako Pure Chemical Corporation, Japan).

### Stable isotope labeling to the calluses

The medium was prepared for the growth of the calluses as following: Murashige Skoog salt (1.38 g) and vitamin (1000×, 300 μL), sucrose (9 g), 1-naphthaleneacetic acid, and kinetin (5 μM final concentration), and gelrite (0.2%) in water (300 mL). The pH was adjusted to 5.7 with KOH. The solution was then autoclaved for 15 min at 121°C.

A medium aqa. solution was autoclaved (3 mL) and added to a well of a 6-well plate (BMBio, Japan). Then, 30 μL of 100 mM aqa. solution of ^13^C-labeled amino acid and its non-labeled counterpart were filtered for sterilization, followed by the addition to the medium to achieve a final concentration of 1 mM. A piece of the callus (5-by-5 mm size) was placed in the center of the well. The plate was incubated at 23 °C in a dark growth chamber. The medium was refreshed once a month. Calluses were harvested after four months. Three biological replicates were prepared for all samples.

### Metabolite extraction and untargeted LC-MS/MS analysis

Freeze-dried samples were extracted and analyzed as described in the previous study^3^.

### S-plot analysis

SIMCA-P (v 12.0.1) was used in this study. Pareto scaling was applied with the default parameters.

### Data processing for metabolome network analysis

Data matrix was created using MassLyncs 4.2. In preliminary processing for metabolome network analysis, isotopic ions in regions (monoisotopic ions (M) + 1.0034/2.0068 ± 0.01 Da) were removed to simplify MS/MS spectra in ^13^C-labeled and non-labeled data. The scores of MS/MS similarity by dot products were calculated for the simplified ^13^C-labeled and non-labeled MS/MS spectra under the following conditions: retention time ±0.05; difference of *m/z* value on precursor ion, 0 ≤ n ≤ 5; number of production ≥ 3. The calculation was performed according to the previous study^3^. Nodes representing ions were clustered using the spinglass method in R (https://igraph.org/r/doc/cluster_spinglass.html). The parameters on gamma and spins were changed to 1.5 and 200, respectively. The information of nodes and edges on the metabolome network analysis is available at DROP Met (http://prime.psc.riken.jp/menta.cgi/prime/drop_index). Metabolome networks using the similarity scores obtained by dot products were visualized using the PlaSMA database (http://plasma.riken.jp/).

### Orally feeding hypertensive mice with asparaptine A

Experiments were outsourced to a company (UNITECH Co., Ltd., Japan). These experiments were conducted in accordance with the regulations of the Act on Welfare and Management of Animals, the Standards relating to the Care and Keeping and Reducing Pain of Laboratory Animals, the Basic Guidelines of the Ministry of Education, Culture, Sports, Science and Technology, the Guidelines for Proper Conduct of Animal Experiments, and the Standards relating to the Methods of Destruction of Animals. In addition, the implementation of this study has been approved by the Animal Experiment Committee of UNITEC Corporation (AGR RK-171011-100).

A total of 14 mice (hRN8-12 × hAG2-5, F1 generation) were moved from a rearing place to a site for measuring their blood pressure. The animals were reared for 6–8 weeks for adaptation to the following conditions: room temperature 22°C-26°C; humidity 40%-65%; light 8:00 AM-20:00 PM, cage, each per mouse; cage exchange or cleanup, every week; and feed and water, discretionary. Asparaptine aqa. solution (2.5 mg/mL) was orally fed to seven mice (age: 15-16 weeks) for 50 μg/g body weight, and water was fed to another seven mice as a control group. Blood pressure was measured by the tail-cuff method at one, two, and three hours and two days after feeding.

## Results and Discussion

The biosynthetic pathway of asparaptine (called asparaptine A hereafter) is largely unknown; however, it has been suggested that the asparagusyl moiety of asparaptine A may be biosynthesized from the amino acid valine. A few steps from valine were only revealed with radio isotope labeling^15^. To understand whether valine can be used as a starting material for the asparagusyl moiety, stable isotope labeling was performed [**Supporting Information (SI) Figure S1**]. Since asparagus is a perennial plant, this makes it difficult to use the asparagus plant itself for stable isotope labeling. In this study, a callus line derived from green asparagus^16^ was used instead.

To increase the labeling rate, pieces of asparagus callus were grown for four months in medium containing ^13^C-labeled or non-labeled valine, which were then harvested for untargeted analysis. Comprehensive data acquisition was performed using liquid chromatography-tandem mass spectrometry (LC-MS/MS). All scanned MS/MS data were then output using MassLyncs. To evaluate the quantity of metabolites that were labeled with ^13^C valine, tracing of endogeneous ^13^C-labeled and non-labeled valine was performed (**Figure 1**), which indicated that the labeling was successful and the average labeling rate was approximately 60% in three biological samples.

**Figure 1.**
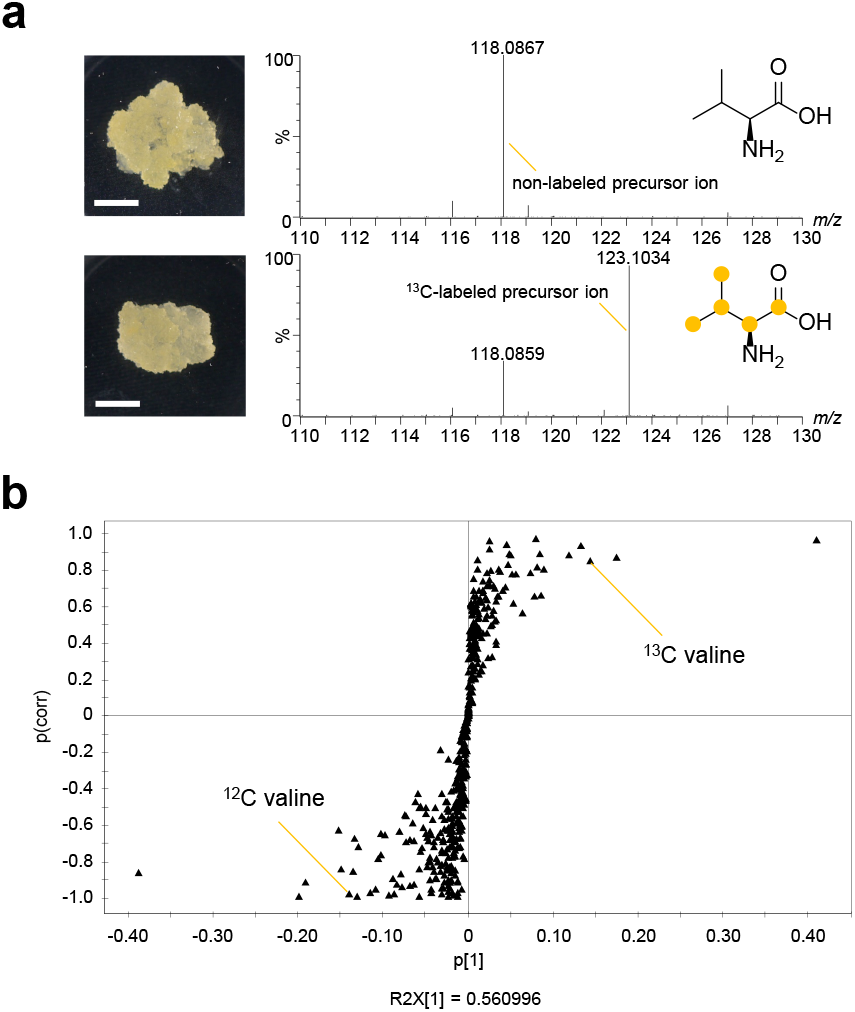
^13^C labeling of asparagus calluses. (a) Representative MS spectrum of endogenous valine in ^13^C-labeled and non-labeled calluses. (b) S-plot using ^13^C-labeled and non-labeled metabolome data. Differences indicated presence of ^13^C-labeled and non-labeled valine-derived metabolites. Scale bar, 0.5 cm.

Using the MS/MS spectra, similarity scores were calculated using dot product method. A metabolome network using both ^13^C-labeled and non-labeled data was created with the following conditions (similarity score ≥ 0.8; mass range, *m/z* 100–500; difference of *m/z* value on precursor ion 1 ≤ n ≤ 5) (**Figure 2; SI Figure S2**). Each node (ion) is represented in a community that consists of other nodes. The communities are constructed when nodes show high similarity scores to each other. In this analysis, the nodes from ^13^C-labeled and non-labeled metabolites were connected with the MS/MS similarity score >0.8. The community members appeared to derive from one peak, indicating that multi-scanned MS/MS spectra in a metabolite peak were summarized to a single community. Possible intermediates and analogs were searched by their exact mass for [M + H]^+^. Communities that included nodes of asparaptine, nodes of a possible intermediate, and distinct nodes for two possible asparaptine analogs were detected with their exact mass (**SI Figures S3–8**). In this analysis, asparagusic acid and its glucose ester could not be detected in the samples.

**Figure 2.**
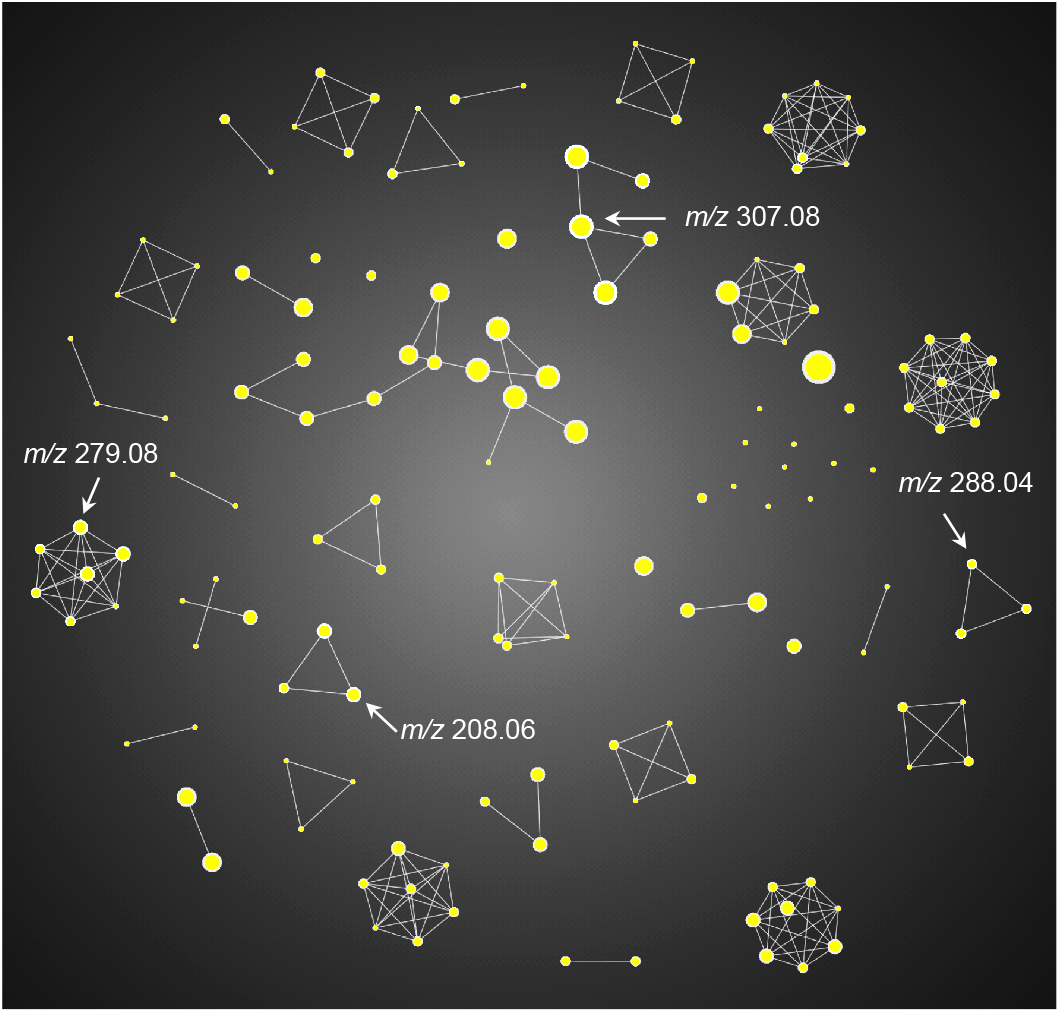
Metabolome network analysis to pair ^13^C-labeled and non-labeled metabolites. The metabolome network was created with the following conditions (similarity score ≥0.8; mass range *m/z* 100–500; difference of *m/z* value on precursor ion 1 ≤ n ≤ 5). Each node indicates a community that consists of nodes (ions), and the edge indicates a similarity score. When nodes have high density, communities were created. When a node in a community share a similarity score to other nodes outside, the community was linked with other communities.

The incorporation of ^13^C to these metabolites was then evaluated by LC-MS/MS. To determine whether the asparagusyl moiety is derived from valine, comparative analysis of ^13^C-labeled and non-labeled MS/MS spectrum was performed (**Figure 3a**). The ion peak of non-labeled asparaptine A was identified using an authentic standard compound^14^. In the ^13^C-labeled MS/MS spectrum, mass-shifted product ions were confirmed. The fragmentation patterns indicated that asparagusyl moiety of asparaptine was derived from valine. Comparative analysis on the possible intermediate was also performed (**Figure 3b**), which showed the exact mass of *S*-(2-carboxy-*n*-propyl)-L-cysteine as [M + H]^+^. A previous study suggested that the S atom is derived from the attachment of the cysteine moiety^15^. The precursor ion observed at *m/z* 208.0646 was found to be nearly identical to the theoretical one (*m/z* 208.0644 for C7H14NO4S [M + H]^+^). The fragmentation patterns indicated that the bonds of N and S were cleaved in the cysteine moiety, suggesting that this metabolite is derived from the cysteine derivative.

**Figure 3.**
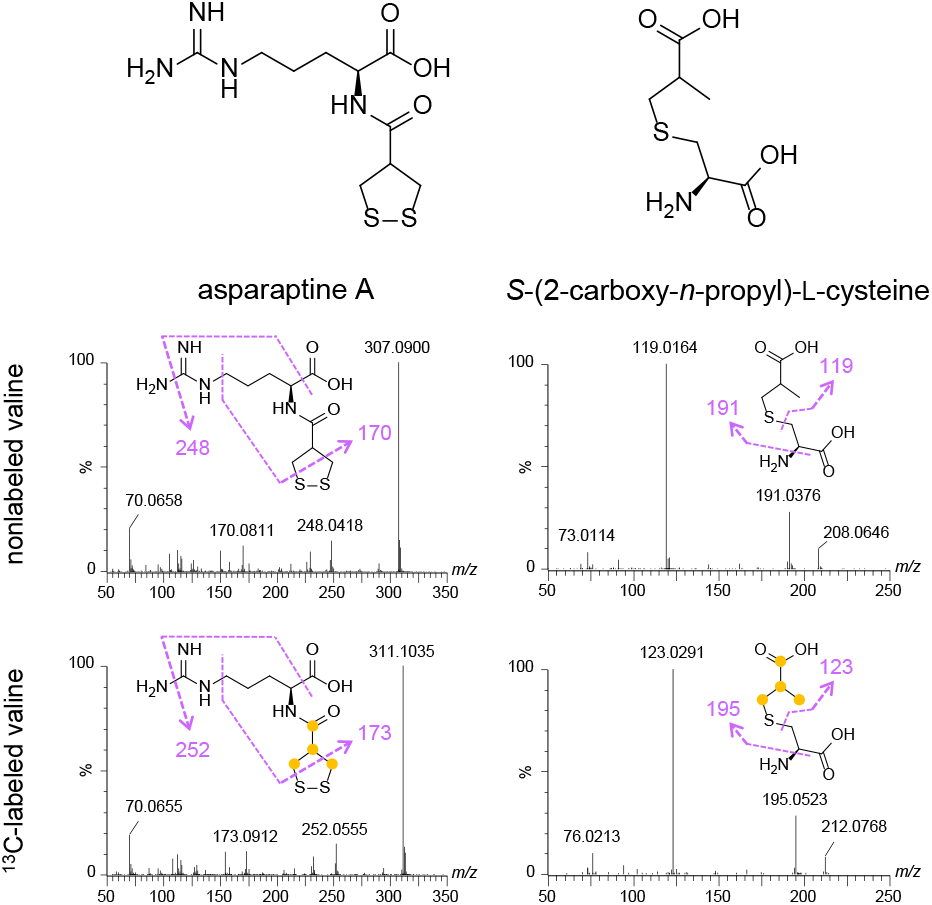
Structural analysis of asparaptine A and *S*-(2-carboxy-*n*-propyl)-L-cysteine. Left panel. Incorporation of ^13^C to the asparagusyl moiety in asparaptine A. This non-labeled asparaptine A was identified using an authentic standard compound [Level 1, the guideline of Metabolomics Standards Initiative (MSI)]. Right panel. Incorporation of ^13^C to the 2-carboxy-*n*-propyl moiety in *S*-(2-carboxy-*n*-propyl)-L-cysteine (Level 3, the guideline of MSI). On the basis of paired ^13^C-labeled and non-labeled MS/MS spectra, MS/MS spectra were demonstrated using MassLyncs.

We hypothesized that, of the two possible analogs, one may be a conjugate of lysine and asparagusic acid and the other one may be a conjugate of histidine and asparagusic acid (**SI Figures S4**). To elucidate their structure, the MS/MS spectrum acquired from calluses labeled with ^13^C valine and ^13^C lysine/^13^C histidine were compared for fragmentation analyses (**Figure 4**). The analysis revealed the incorporation of ^13^C valine to the asparagusyl moiety and that of ^13^C-lysine/^13^C histidine to the lysyl or histidyl moiety. The fragmentation patterns identified the cleavage of amino and carboxyl moieties, showing that the asparagusyl moiety is conjugated to the amino moiety at the α position of lysine/histidine. Thus, asparaptine was renamed as asparaptine A. The lysine- and histidine-type analogs were named as asparaptines B and C, respectively. The existence of these analogs suggest that additional asparaptine analogs may also exist in asparagus plants or calluses. A cutting-edge LC-MS/MS instrument with higher sensitivity is expected to detect the additional analogs.

**Figure 4.**
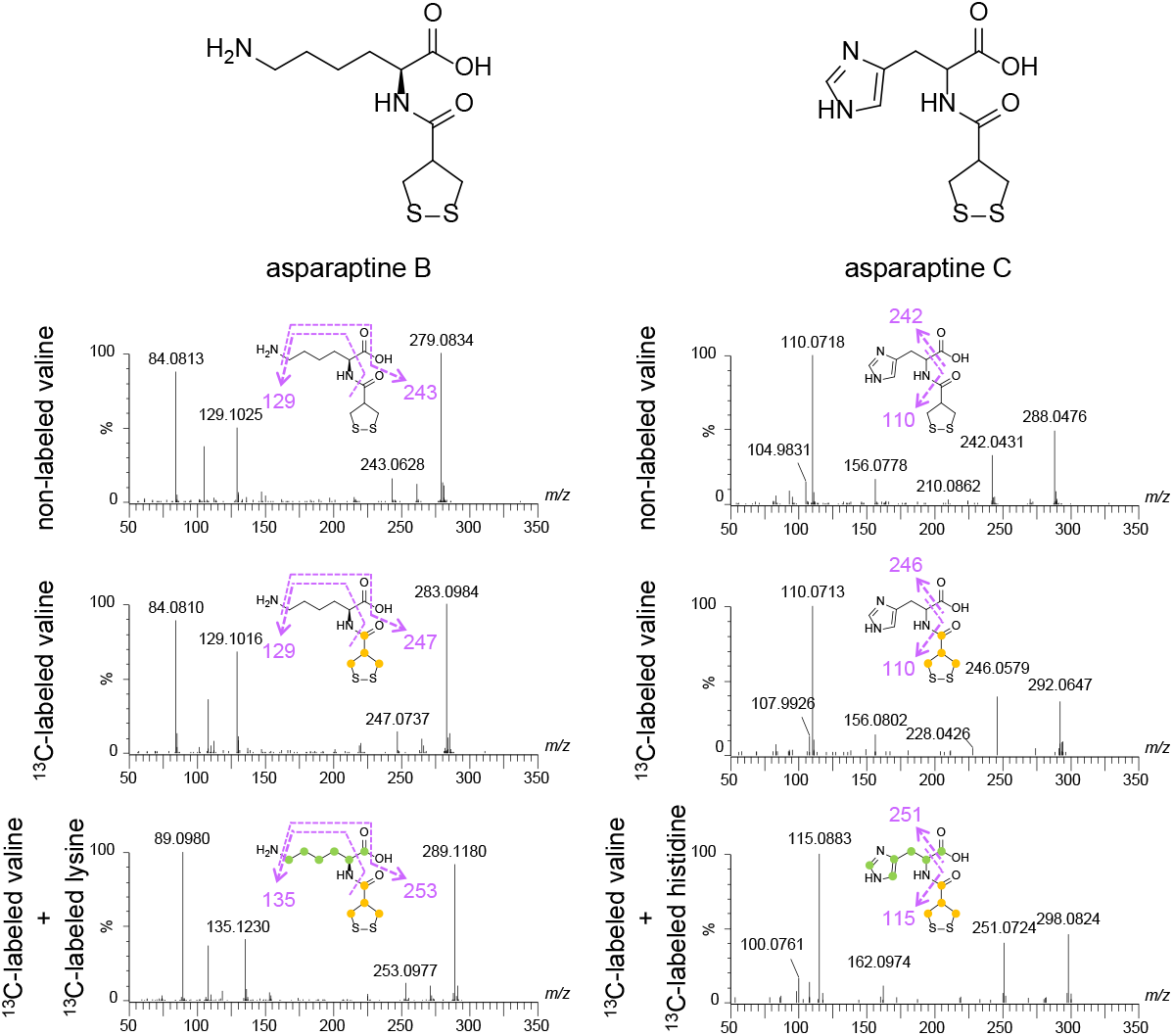
Structural analysis of asparaptines B and C. In addition to using paired ^13^C-labeled and non-labeled MS/MS spectra, a doubly labeled MS/MS spectrum was employed for this analysis. Top. MS/MS spectrum of non-labeled asparaptines B and C. Middle. MS/MS spectrum of asparaptines B and C labeled with ^13^C valine. Bottom. MS/MS spectrum of asparaptines B and C labeled with ^13^C-labeled valine and ^13^C-labeled lysine/histidine. On the basis of paired ^13^C-labeled and non-labeled MS/MS spectra, MS/MS spectra were demonstrated using MassLyncs.

As mention above, asparaptine A shows the ACE inhibitory activity *in vitro.* Interestingly, the conjugate of proline and asparagusic acid [named asparaptine K in this study (**SI Figure S4**)] has been reported as a blood pressure-lowering agent^17^. To confirm blood pressure-lowering effect *in vivo,* hypertensive mice were orally fed with asparaptine A (**Figure 5**). Both the systolic and diastolic blood pressure in the asparaptine A-fed mice remarkably decreased to 20 mmHg lower than those in control group in two hours after feeding (**Figures 5a** and **5b**). After two days, the blood pressure completely returned to normal. This observation suggested that asparaptine A has a rapid action in reducing blood pressure. Moreover, the heart rate in the asparaptine A-fed mice significantly decreased in two hours, before recovering quickly (**Figure 5c**). These results suggest that asparaptine A may not only have effects as an ACE inhibitor, but also has β-antagonistic effects. These effects are well-known to be preventive for cardiovascular diseases^18,19^. Short-acting ACE inhibitors with strong anti-hypertensive effects are useful for patients with aortic dissection or rupture in emergency department. However, most ACE inhibitors are well-known as long-acting anti-hypertensive agents, which makes them difficult to use for patients who require urgent surgical intervention. Asparaptine A may be an important candidate as an anti-hypertensive drug in future medical use due to the strong observed anti-hypertensive effects combined with rapid degradation in the mice.

**Figure 5.**
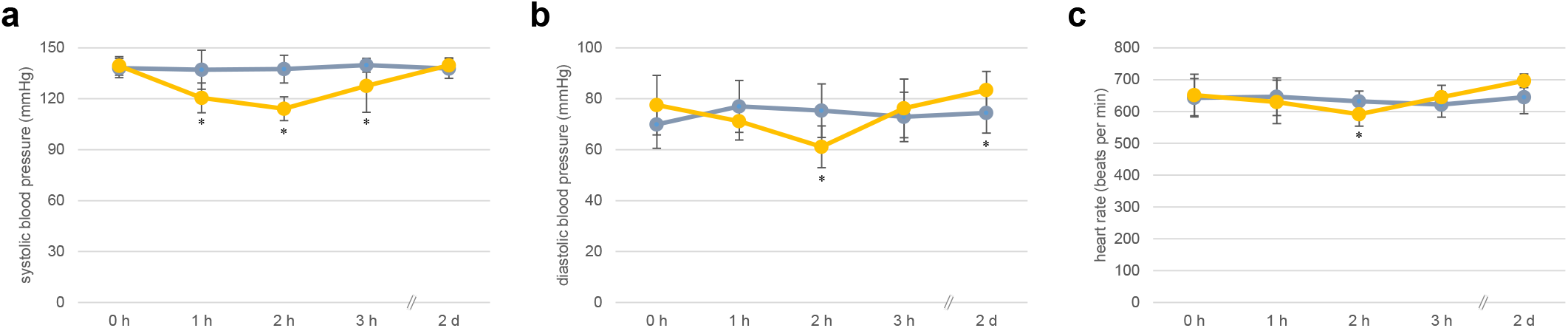
Effect of asparaptine A on hypertensive mice. (a) Systolic blood pressure. (b) Diastolic blood pressure. (c) Heart rate. Blue line indicates the water-fed group (control), and yellow line indicates the asparaptine A-fed group. Standard deviations (error bars) were calculated from the results of seven biological replicates. Asterisk indicates significant differences between the groups at the annotated time point (Student *t*-test, *p* < 0.05).

## Conclusion

In this study, we performed the MS/MS similarity-based network approach to analyze ^13^C-labeled and non-labeled metabolome data from asparagus calluses. The analysis characterized metabolite pairs despite differences in the *m/z* value. Four paired ions were analyzed to determine the structure of an unknown pathway intermediate *S*-(2-carboxy-*n*-propyl)-L-cysteine, asparaptine A, and two previously unidentified analogs asparaptines B and C. This analysis enables the automatic extraction of accurate pairs of ions derived from ^13^C-labeled and non-labeled metabolites in metabolomics data. The analysis is applicable to software programs that require off-the-shelf approaches to pair stable isotope-labeled and non-labeled metabolites.

## Associated Contents

### Supporting Information

The Supporting Information is available free of charge at online.

**Figure S1.** ^13^C labeling of asparagus calluses.

**Figure S2.** Parameters of the metabolome network analysis.

**Figure S3.** Searched metabolites in the metabolome network analysis.

**Figure S4.** Searched possible analogs of asparaptine A.

**Figure S5.** Linking nodes (^13^C-labeled) to their counterparts (non-labeled) on asparaptine A.

**Figure S6.** Linking nodes (^13^C-labeled) to their counterparts (non-labeled) on *S*-(2-carboxy-*n*-propyl)-L-cysteine.

**Figure S7.** Linking nodes (^13^C-labeled) to their counterparts (non-labeled) on asparaptine B.

**Figure S8.** Linking nodes (^13^C-labeled) to their counterparts (non-labeled) on asparaptine C.

### Author Information

Yutaka Yamada

RIKEN Center for Sustainable Resource Science, Yokohama 230-0045, Japan

Tomoko Nishizawa

RIKEN Center for Sustainable Resource Science, Yokohama 230-0045, Japan

Tetsuya Mori

RIKEN Center for Sustainable Resource Science, Yokohama 230-0045, Japan

Takashi Asano

Iwate Medical University, Iwate 028-3694, Japan

Masanari Kuwabara

Toranomon Hospital, Tokyo 105-8470, Japan

Kazuki Saito

RIKEN Center for Sustainable Resource Science, Yokohama 230-0045, Japan

### Author Contributions

R.N. designed the research. R.N., T.N., and T.A. prepared the callus samples. R.N. and T.M. acquired the metabolome data. R.N. and Y.Y. performed the metabolome network analysis. R.N. analyzed all the data. M.K analyzed the data of blood pressure. R.N. discussed the research with all the co-authors. R.N. wrote the manuscript.

## Acknowledgments

We would like to thank Hiroshi Tsugawa for technical advices (RIKEN CSRS) and Kouji Takano (RIKEN CSRS) for technical assistance. The hypertensive mice were provided by the RIKEN BRC through the National Bio-Resource Project of the MEXT, Japan.

## Notes

The authors declare no competing financial interest.

