## Supplementary material for "Tandem mass spectrum similarity-based network analysis using ^13^C-labeled and non-labeled metabolome data to identify the biosynthesis pathway of the blood pressure-lowering asparagus metabolite asparaptine A": Figures S1 to S8

### SI Figure S1

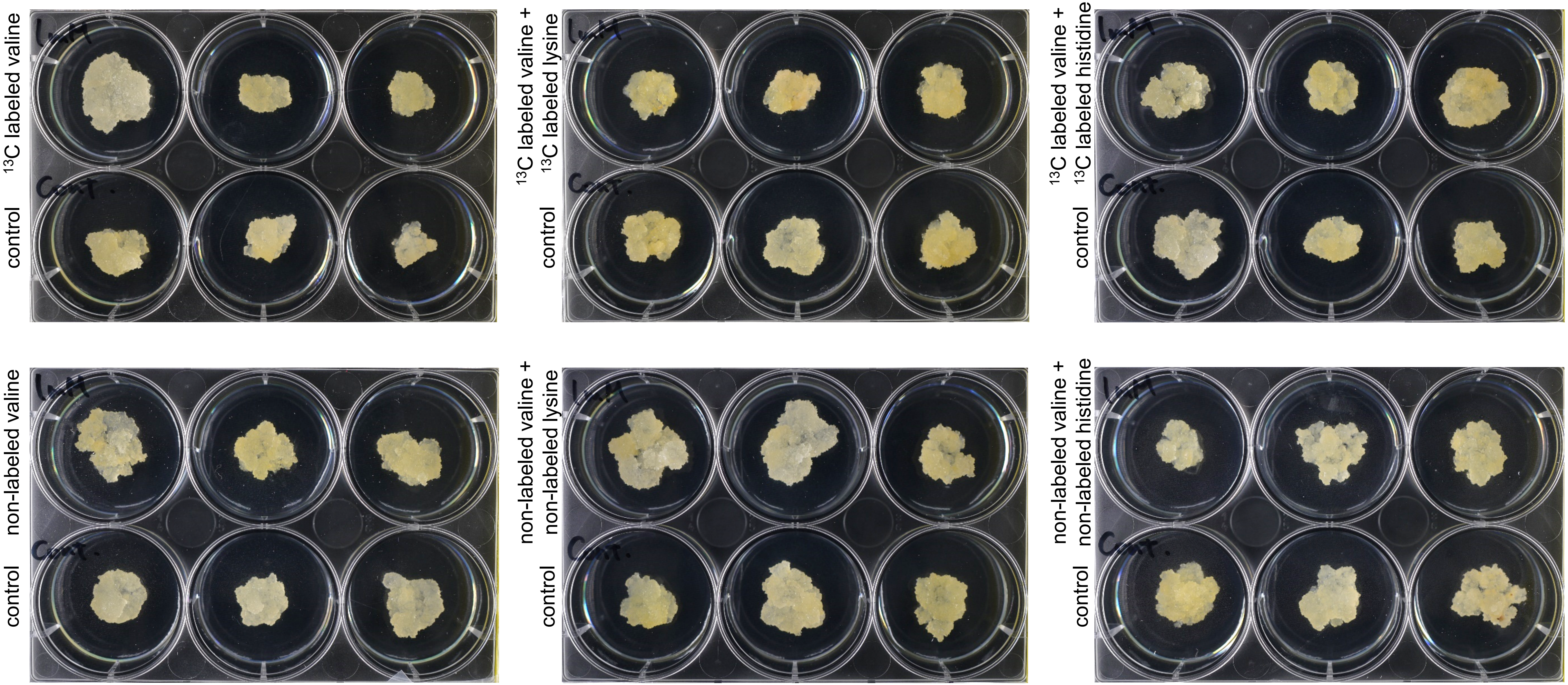


**Figure S1.** $^{13}$C labeling of asparagus calluses.
Amino acid (30 μL of 100 mM stock solution) was added to the agar medium for final concentration (1 mM) . Water (30 μL) was added for control.

### SI Figure S2

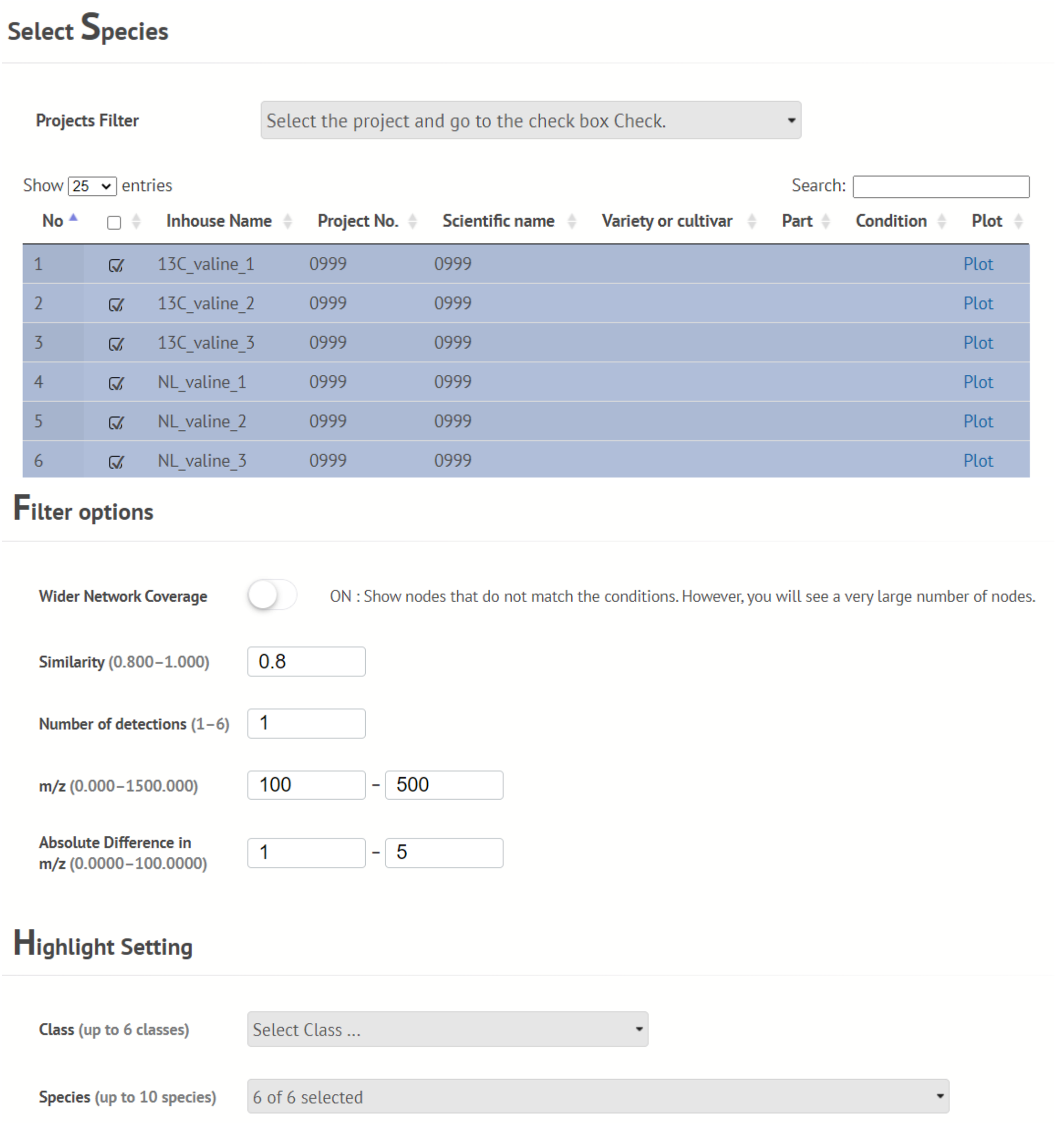


**Figure S2.** Parameters of the metabolome network analysis.

### SI Figure S3

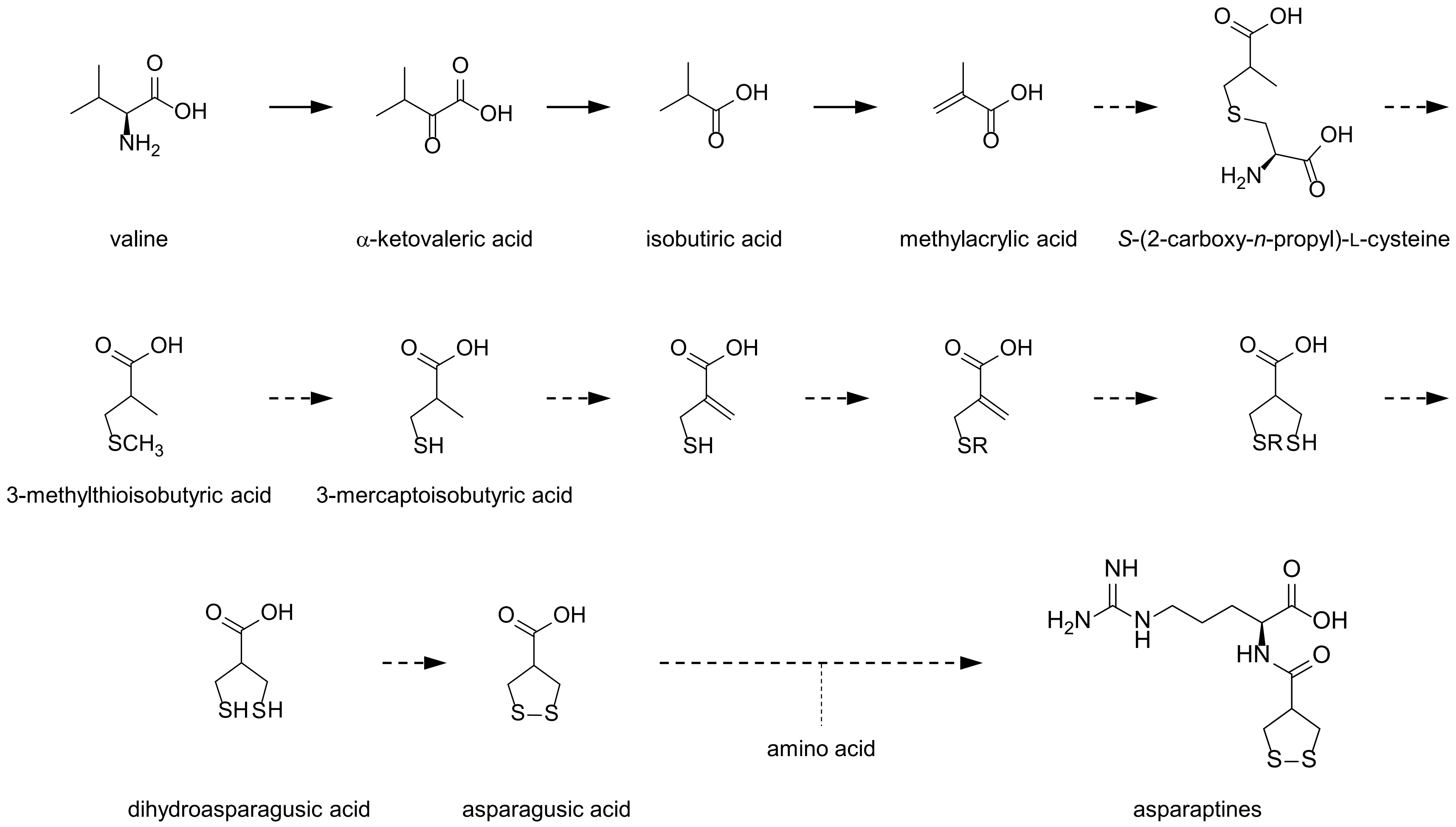


Mitchell and Waring, Phytochemistry (2014), modified
Yanagisawa and Egami, J. Biol. Chem. (1976)

**Figure S3.** Searched metabolites in the metabolome network analysis.

### SI Figure S4

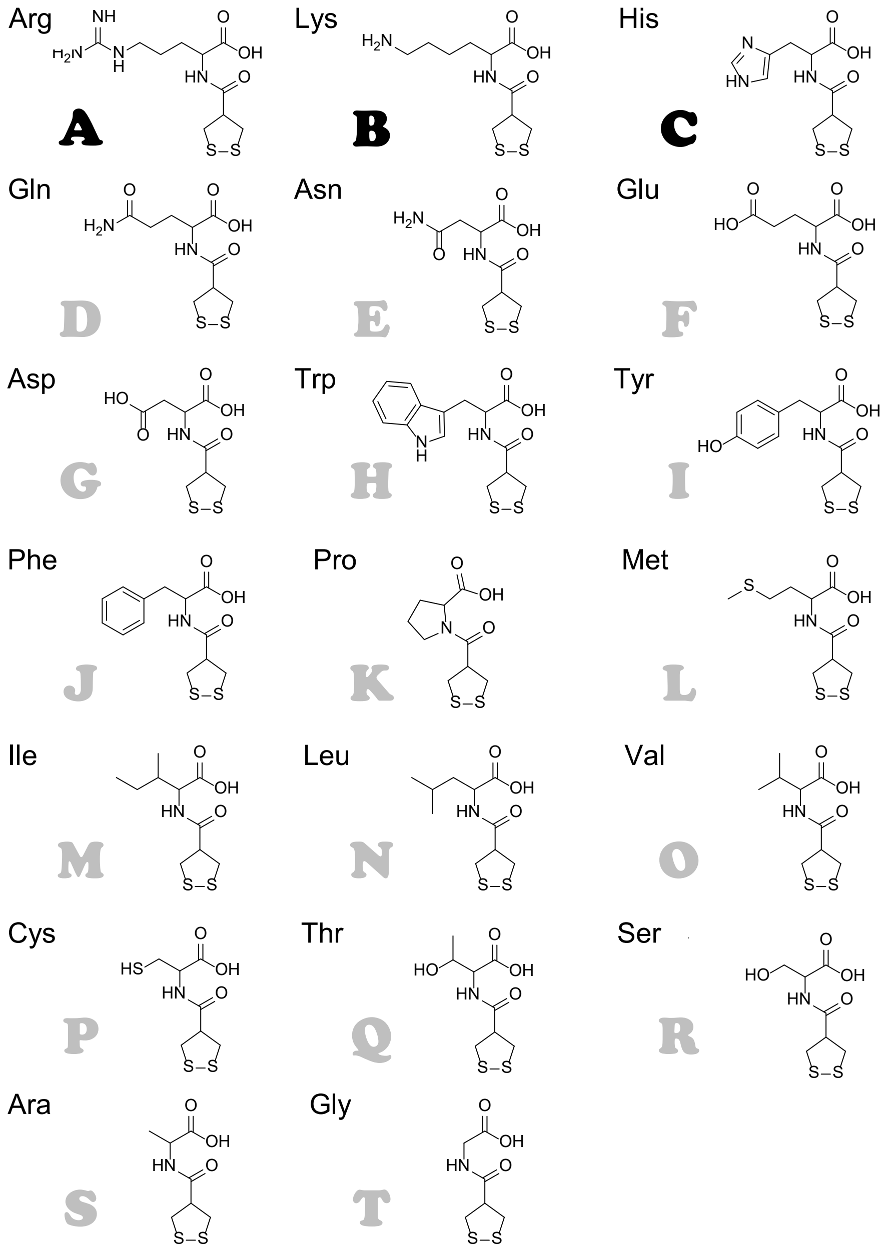


**Figure S4.** Searched possible analogs of asparaptine A.

### SI Figure S5

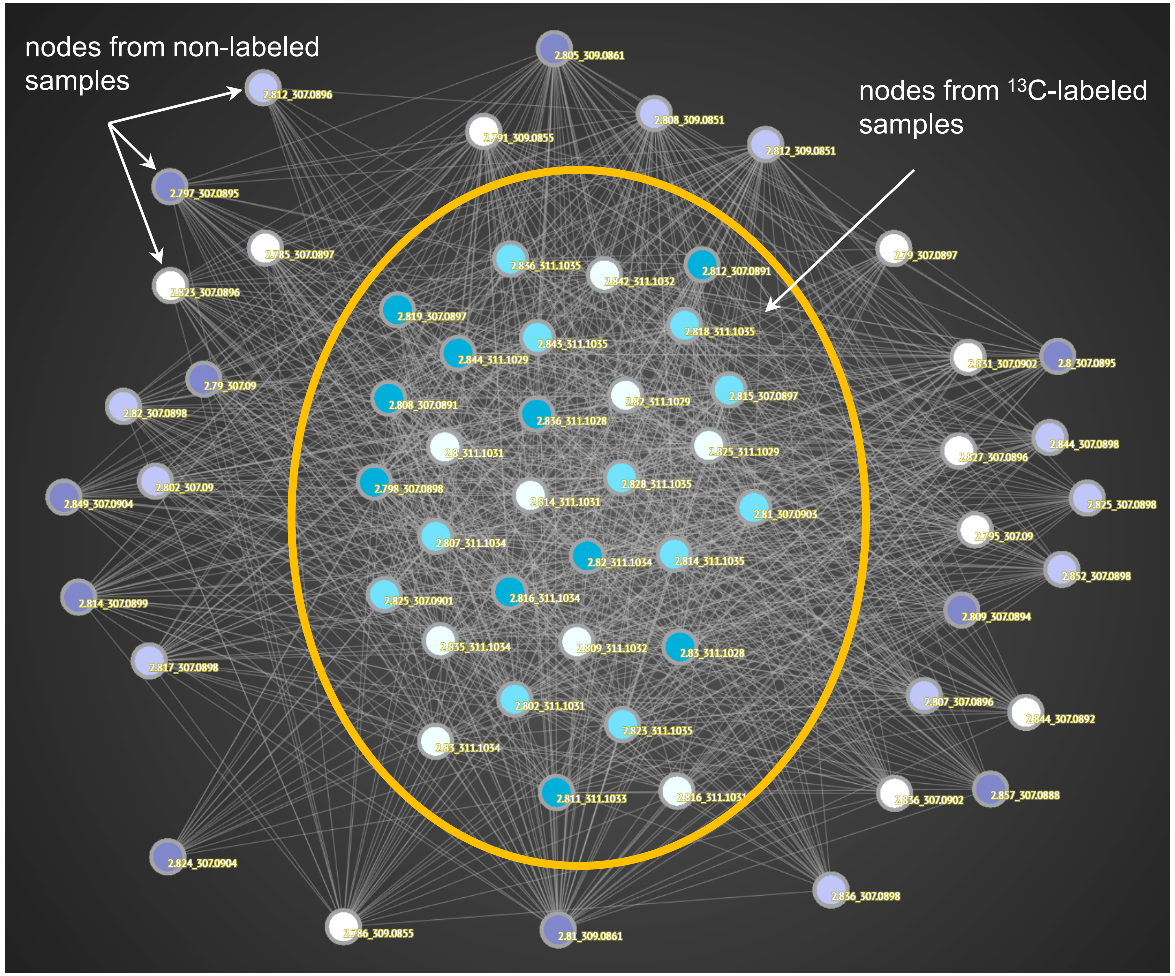


**Figure S5.** Linking nodes ($^{13}$C-labeled) to their counterparts (nonlabeled) on asparaptine A .

### SI Figure S6

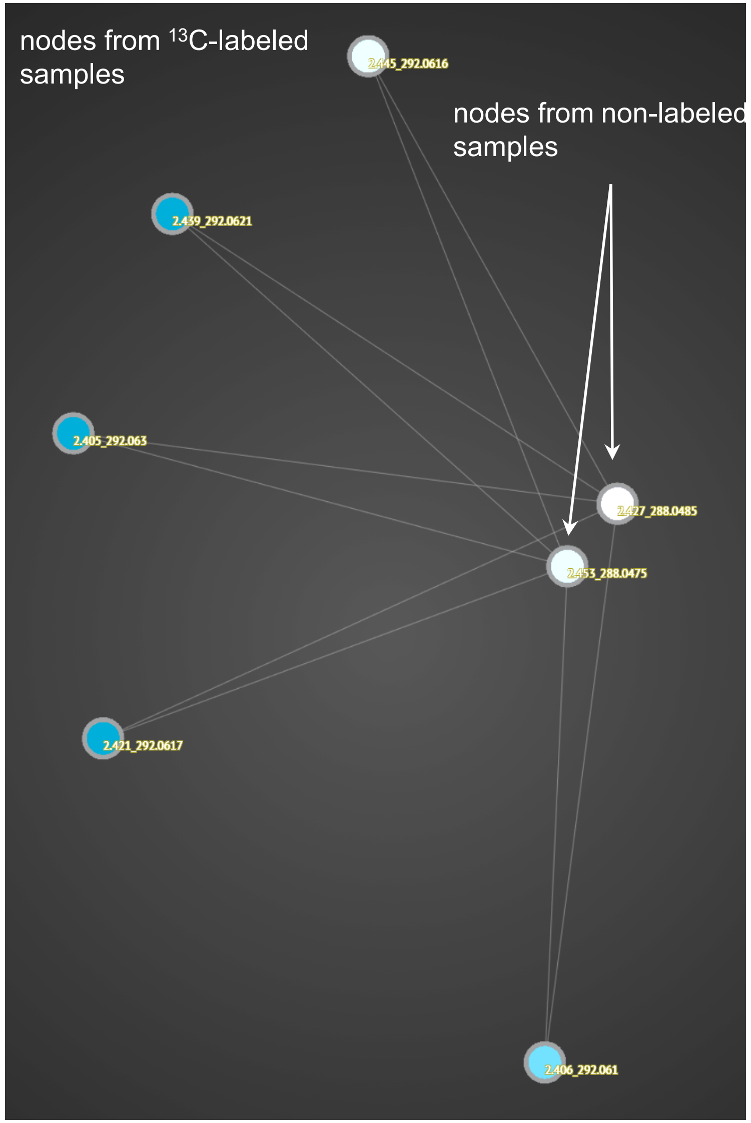


**Figure S6.** Linking nodes ($^{13}$C-labeled) to their counterparts (nonlabeled) on *S*-(2-carboxy-*n*-propyl)-L-cysteine.

### SI Figure S7

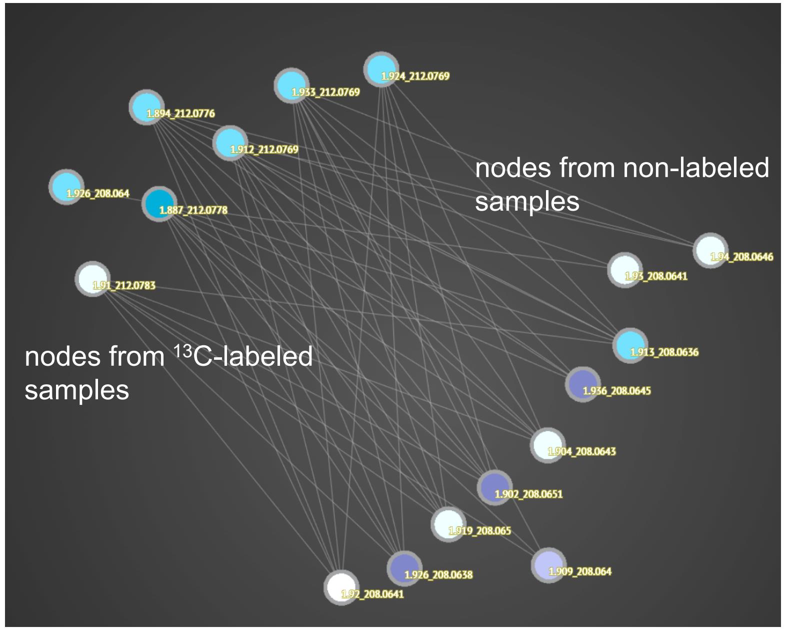


**Figure S7.** Linking nodes ($^{13}$C-labeled) to their counterparts (non-labeled) on asparaptine B.

### SI Figure S8

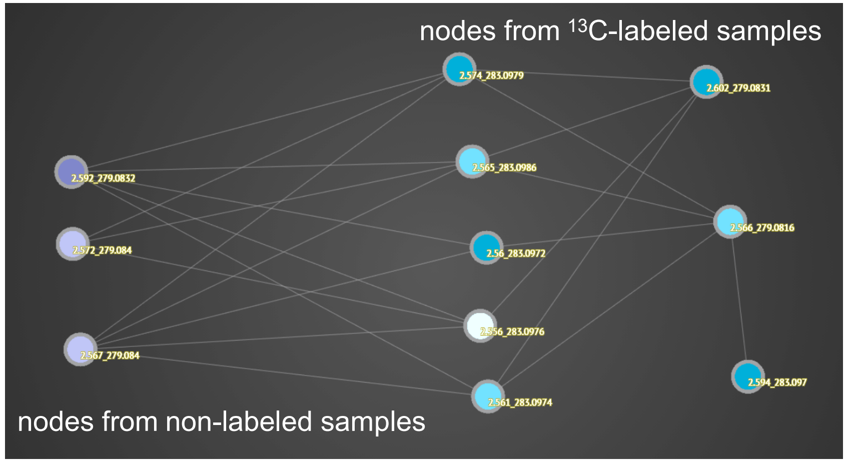


**Figure S8.** Linking nodes ($^{13}$C-labeled) to their counterparts (non-labeled) on asparaptine C.
